# Measuring mosquito-borne viral suitability in Myanmar and implications for local Zika virus transmission

**DOI:** 10.1101/231373

**Authors:** PN Perez, U Obolski, LCJ Alcantara, M Maia de Lima, EA Ashley, F Smithuis, P Horby, RJ Maude, L Zaw, AM Kyaw, J Lourenço

## Abstract

INTRODUCTION: In South East Asia, mosquito-borne viruses (MBVs) have long been a cause of high disease burden and significant economic costs. While in some SEA countries the epidemiology of MBVs is spatio-temporally well characterised and understood, in others such as Myanmar our understanding is largely incomplete. MATERIALS AND METHODS: Here, we use a simple mathematical approach to estimate a climate-driven suitability index aiming to better characterise the intrinsic, spatio-temporal potential of MBVs in Myanmar. RESULTS: Results show that the timing and amplitude of the natural oscillations of our suitability index are highly informative for the temporal patterns of DENV case counts at the country level, and a mosquito-abundance measure at a city level. When projected at fine spatial scales, the suitability index suggests that the time period of highest MBV transmission potential is between June and October independently of geographical location. Higher potential is nonetheless found along the middle axis of the country and in particular in the southern corridor of international borders with Thailand. DISCUSSION: This research complements and expands our current understanding of MBV transmission potential in Myanmar, by identifying key spatial heterogeneities and temporal windows of importance for surveillance and control. We discuss our findings in the context of Zika virus given its recent worldwide emergence, public health impact, and current lack of information on its epidemiology and transmission potential in Myanmar. The proposed suitability index here demonstrated is applicable to other regions of the world for which surveillance data is missing, either due to lack of resources or absence of an MBV of interest.

## Introduction

Common mosquito-borne viruses (MBVs) of global health concern include the dengue (DENV), chikungunya (CHIKV), Zika (ZIKV), yellow fever (YFV), Rift Valley fever (RVFV), West-Nile (WNV) and Japanese encephalitis (JEV) viruses. Due to ongoing globalization and climatic trends that favour the establishment of vectors and movement of infectious hosts, these pathogens are becoming increasingly detrimental for human public health [1]–[3].

The evolutionary and host-pathogen history of MBVs is vastly diverse, but the population biology of these viruses shares one unifying characteristic: their epidemiological dynamics and epidemic behaviour is inherently linked to the underlying population dynamics of their vector-species. Mosquito-population dynamics are known to be dictated by a wide range of factors, such as climate, altitude, population density of humans or other animals, air or waste pollution levels, and natural or artificial water reservoirs [4]–[7]. While most of these factors can dictate absolute population sizes (carrying capacity), seasonal oscillations are largely driven by natural climate variations. South East Asia (SEA) is the most densely populated region of the world and has experienced rapid urbanization in the past century. Such demographic factors and tropical climate are believed be the main drivers of the success of *Aedes* mosquitoes in the region, the main genus for the transmission of MBVs such as DENV, CHIKV and ZIKV. Historically, surveillance of MBVs has been highly heterogeneous in the region. For instance, a few countries such as Vietnam, Thailand and Singapore often serve as global references for DENV epidemiology, reporting spatiotemporal epidemiological and genetic data spanning several decades [8], [9]. Other countries, such as Myanmar, report epidemiological data at lower spatio-temporal resolutions [10]–[12]. For instance, published DENV case counts over time are reported at national level with total counts per year at the state level, but spatio-temporal data at the district or lower levels is not freely available. In the context of countries with incomplete data coverage for a robust understanding of the local epidemiological determinants and epidemic potential of MBVs, mathematical frameworks, and in particular dynamic models, are essential to close the gaps in knowledge.

Recently, we have developed and applied a climate-driven mathematical framework to study epidemics of MBVs such as DENV [13], [14] and ZIKV [15]. The success of this framework stems from its data-driven approach, including temporal climatic series used to parameterise both vector and viral variables under mathematical relationships derived from experimental studies. The reliance on local climatic variables as its main input makes the framework general enough to be applied to different geo-locations. Here we translate our experience with this framework to introduce an MBV suitability index, which we apply to the context of Myanmar. With it, we estimate spatio-temporal patterns of suitability across the country, validating it against existing but incomplete data. Our results contribute to a better understanding of Myanmar’s MBV transmission potential in both time and space. We discuss the public health and control implications of our findings both generally for MBVs and in particular for ZIKV.

## Methods

Our approach develops from a climate-driven, mosquito-borne mathematical model of viral transmission that has been successfully applied to three MBV epidemics: for the 2012 dengue serotype 1 outbreak in the island of Madeira (Portugal) [13], the 2014 dengue serotype 4 outbreak in Rio de Janeiro (Brazil) [14] and the 2015-2017 ZIKV outbreak in Feira de Santana (Brazil) [15]. In these case studies, given the availability of reported epidemic curves at appropriate spatio-temporal scales, deterministic simulations were used with a Markov chain Monte Carlo fitting approach to derive key local eco-epidemiological parameters, allowing of the estimation of the basic reproductive number (R0) and effective reproductive number (Re).

In the context of Myanmar and the lack of reported case counts at the subnational level, we addressed the potential for MBVs transmission by focusing on the model’s equation for R0 [15], a similar starting point of a recent and successful strategy implemented in the context of YFV in the African continent [16]. R0 is the sum of the transmission potential of each adult female mosquito per human, across the total number of female mosquitoes per human (M), in a totally susceptible human population. Hence, R0 can be expressed as the product of M with each individual mosquito transmission potential P(u,t):

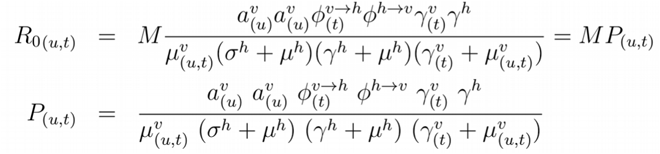

The R0 expression is dependent on time-varying humidity (u) and temperature (t); M is the number of adult female mosquitoes per human (V/N, with N the human population size and V the adult female mosquito population size); a^v^(u) is the mosquito biting rate, dependent on humidity; ϕ^v-h^(t) is the probability of transmission of the virus by infected-mosquito to human, per bite, dependent on temperature; ϕ^h-v^ is the probability of transmission from infected-human to mosquito, per bite; 1/Y^v^(t) is the extrinsic incubation period, dependent on temperature; 1/Y^h^ is the intrinsic incubation period; 1/μ^v^(u,t) is the mosquito life-span, dependent on humidity and temperature; 1/σ^h^ is the human infectious period; and 1/μ^h^ is the human life-span. Each of the climate dependent functions was previously determined by laboratory estimates of entomological data (equations 20-37, [15]).

The individual-mosquito transmission potential P(u,t) is a positive number, with P>1 indicating the capacity of a single adult female mosquito to contribute to epidemic expansion, and P<1 indicating otherwise. However, this threshold around 1 does not necessarily equate to the classic epidemic threshold of R0>1 (or Re>1), since P(u,t) critically ignores the total number of female mosquitoes per human (M). In other words, an epidemic threshold may be reached, for instance, with P<1 if M>>1. Here, we argue and demonstrate that P(u,t) holds critical information on transmission seasonality (timing) and amplitude (relative epidemic potential between seasons and regions), both driven by the inherent climatic variables affecting viral and entomological factors. We denote P(u,t) as the *mosquito-viral suitability index P*, using a complementary terminology to existing *vector suitability* indices which more generally consider entomological factors and / or vector-population sizes [17]–[19]. As seen above, P(u,t) is a complex expression containing human, entomological and viral factors, and can thus be parametrized for any species of virus, host or vector, and estimated for any region for which humidity and temperature are available.

While some parameters used to calculate P can be quantified through known constant values, others follow mathematical expressions that depend on three scaling factors, α, ρ and η, that are used to modulate the baseline relationships between climatic variables and entomological parameters:

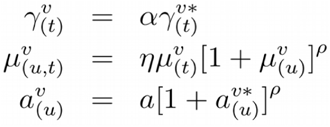

Here, a is the baseline biting rate, and the terms marked with * are the actual functions defined in the empirical studies, which can be found in Materials and Methods of Lourenco et al. [15]. The multiplicative coefficients η and α are used in the temperature dependent components of the adult mosquito mortality and incubation period, respectively. Their inclusion does not alter the relative effect of temperature variation on the entomological parameters *per se*, but allows for the parameter’s baselines to be different from the ideal laboratory conditions of the original research (e.g. [20]). In practice, the effect of temperature on these parameters can be considered to be the same as observed under laboratory conditions if η~1 and α~1, or weaker if <1 or higher if >1. The exponential parameter ρ allows instead to modulate the strength by which adult mosquito mortality and biting rate react to deviations from local mean humidity. In practice, the effect of humidity can be switched off when *ρ* tends to 0 and made stronger when ρ >1. For a discussion on possible biological factors that may justify these factors divergence from 1 please refer to the original description of the method [13] and in a separate study by Brady and colleagues [21].

Fitting exercises to MBV epidemic curves that would allow for quantitative estimations of scaling factors α, ρ and η were not possible for Myanmar due to the lack of reported cases at appropriate spatio-temporal scales. To overcome this limitation, we ran a parameter sweep on the three factors in the range 0-10, and drew the combination of three values that would derive a yearly mean life-span of adult mosquitoes of ~9 days and an extrinsic incubation period of ~5 days. These heuristics are based on prior knowledge for ZIKV and DENV transmission estimations with the same model in three different regions [13]–[15], which are themselves informed by reported biological ranges for *Aedes* mosquitoes [15], [21]–[24]. Please refer to the *Data* section for a description on the epidemiological and climate time series used in this study and the section *Parameters specific to Myanmar* section for all subnational values found and used for the scaling factors α, ρ and η. Constant parameter values used were: human lifespan of 64 years [25], human infectious period of 5.8 days, biting rate of 0.25 per day, human incubation period of 5.9 days and the infected-human to mosquito probability of transmission per bite of 0.5 (as previously modelled [15]).

## Results

We first tested the index P in the context of publicly available DENV case count data at the national-level for the period between 1996 and 2001 [12]. For this, we estimated P for each district using available local climatic data (2015-2016), further aggregating and averaging P across all districts of the country and per month (Figures 1 A1-2). While the epidemiological and climatic data available for the analysis were from different time periods, we found that the estimated P for both 2015 and 2016 presented seasonal fluctuations in sync with mean DENV counts from multiple years (1996-2001). The dynamics of P at the country level further presented key signatures in accordance to Myanmar’s climatic seasons. Namely, (i) a sharp increase in transmission potential during May and June, coincident with the onset of the rainy season (Jun - Oct), and (ii) a trough in potential in the middle of the hot and dry season (Mar - May). A linear correlation between mean DENV counts (1996-2001) and mean index P (2015-2016) showed that ~76% of the variation in case count dynamics could be explained by the index with statistical significance (p-value=2.4×10^−4^, Figure 1 B3).

**Figure 1.**
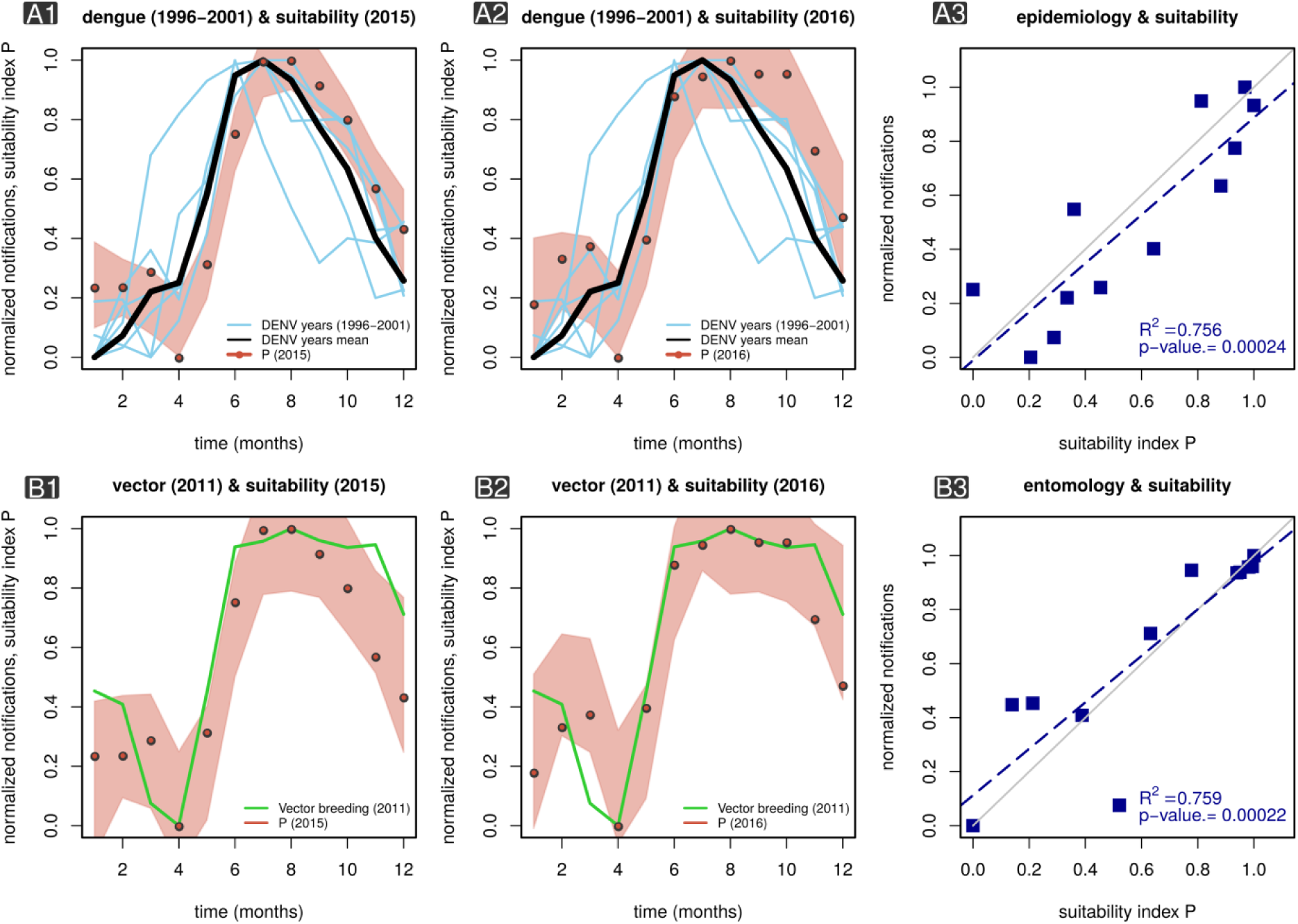
MBV suitability index P and ento-epidemiological tíme series in Myanmar. **Panel A1** presents the mean estimated index P across Myanmar for 2015 (red dots, locally weighted smoothing bounds within red area) superimposed on monthly case counts of DENV for several transmission seasons (1996-2001, blue lines) in Myanmar. The black line is the mean DENV case counts 1996-2001. **Panel A2** is the same as B1 but with P estimated for 2016. **Panel A3** is a linear regression of mean DENV case counts (1996-2001) versus the mean index P (2015-2016) as displayed in panels A1-2. **Panel B1** presents the mean estimated index P in Yangon for 2015 (red dots, locally weighted smoothing bounds within red area) superimposed on monthly number of major breeding containers in 2011 (green lines) for Yangon. **Panel B2** is the same as B1 but with P estimated for 2016. **Panel B3** is a linear regression of major breeding sites (2011) versus the mean index P (2015-2016) as displayed in panels B1-2.

Because we propose the index P as a measure of suitability independent of the total number of (female) mosquitoes, it is critical to have empirical support on whether its oscillatory behaviour (timing and amplitude) correlates with local measures of mosquito population size. Studies including mosquito surveys in Myanmar are scarce in the literature, mostly targeting vectors related to malaria transmission and focusing on their spatial distribution and species diversity (e.g. [26], [27], [28]). Here, we use data collected in a study of *Aedes aegypti* abundance in Yangon for the year of 2011 [29]. As far as we know, this is the only study with larval indices measured monthly over the period of at least one year. We compared the published data with our estimations of index P for Yangon in 2015 and 2016 (Figures 1 B1-2). As previously shown for DENV counts (Figure 1 A1-2), the seasonal fluctuations of index P and larval indices were synchronous in time. A linear correlation between Yangon’s larval indices (2011) and index P (2015) showed that ~76% of the variation in mosquito abundance measure, a proxy for adult mosquito population size, could be explained by the index P with statistical significance (p-value=2.2×10^−4^, Figure 1 C3).

As in other places of the world presenting endemicity for MBVs, Myanmar is likely to present significant heterogeneities between districts (and states) in terms of suitability (transmission potential). Apart from studies reporting differences in the total number of MBV case counts across the country, an assessment of suitability in space has not been previously done for Myanmar. Since our results suggested that the index P contains information on the timing and amplitude of observed DENV counts and mosquito population size (Figure 1), we explored the spatial variation of index P across the country.

We looked at the spatial variation of the index P (2015-2016), focusing on its average within the cool dry (Nov - Feb), hot dry (Mar - May) and wet seasons (Jun - Oct) (Figure 2A), and found significant differences between seasons. Namely, the cool dry season presented a generally lower index P across space, at a time when climatic conditions are expected to be less favorable for the mosquito and therefore for suitability. In contrast, the wet season presented the highest index P across space. Importantly, the latter occurs in the same time period (Jun - Oct) in which DENV counts and mosquito abundance measures also peak. Significant variation within each season was also observed, with the cool and dry season presenting more homogeneous suitability across space (standard deviation, SD=3.86) and the wet season presenting the most heterogeneous suitability (SD=4.17).

**Figure 2.**
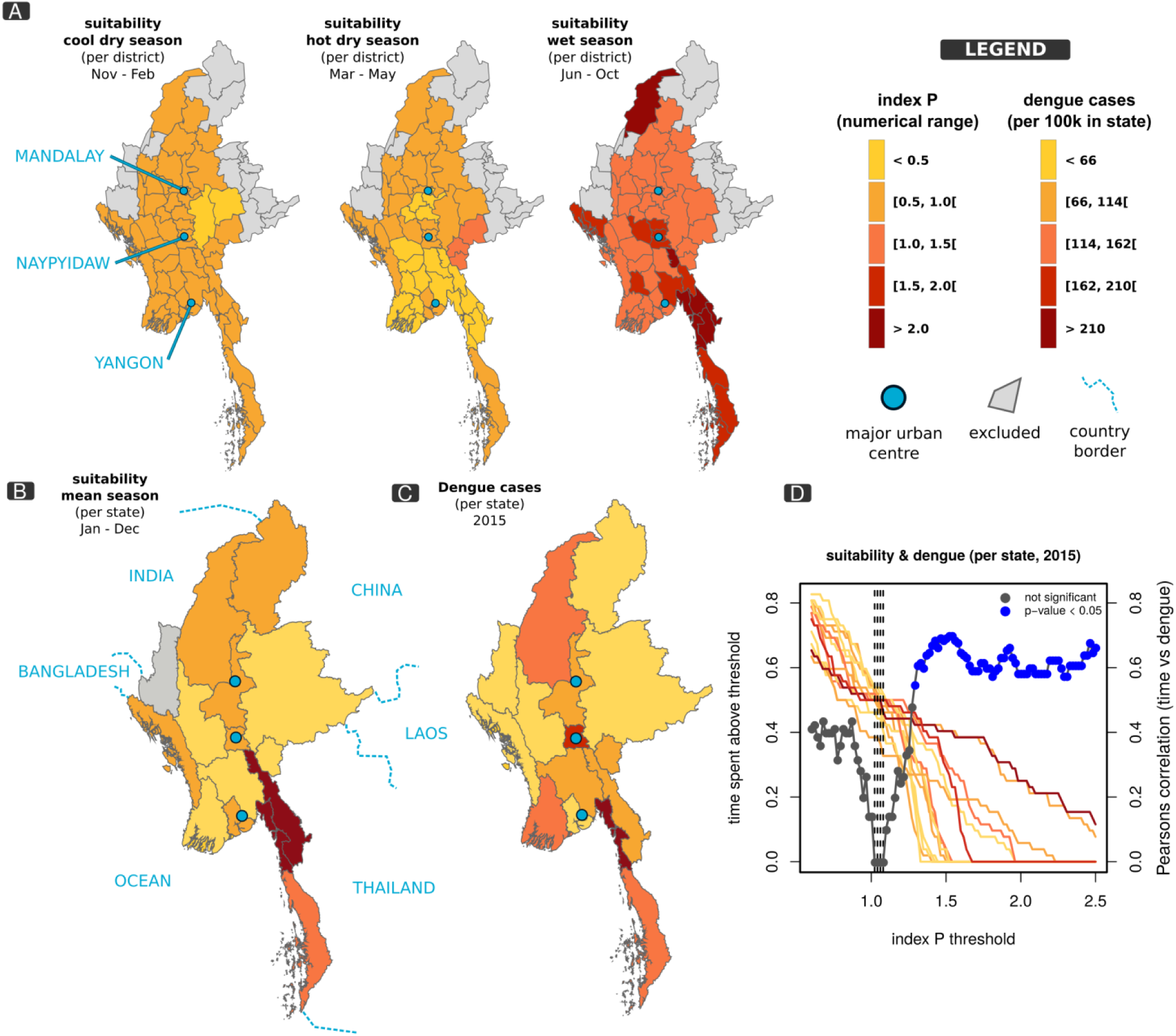
MBV suitability index P and spatial distribution of epidemiological data in Myanmar. **Panel A** shows maps of Myanmar coloured according to mean index P per district in different seasons of the year (as labelled in each map). **Panel B** presents the yearly mean index P per state in 2015 with borders of neighboring countries (named) shown in light blue. **Panel C** presents the number of DENV cases per state in 2015. **Panel D** presents a sensitivity exercise showing the critical index P (~1.5) for which the spatial distributions of dengue cases and mean index P are most correlated in 2015. Colored lines show the amount of time (T) each state spends with index P above a certain threshold (colors related to 2015 DENV case counts, as in map C). Points present Pearson’s correlation coefficient between T and dengue case counts with significant correlations in blue. Dashed vertical lines signal the T values for which the minimum (no) correlation is found. **In all panels**: all model parameters as described in Methods section, except for α, ρ and η as described in section *Parameters specific to Myanmar*.

We next attempted to correlate the spatial distribution of suitability with local mosquito abundance measures and MBV case counts. However, we found no data with spatial resolution for abundance, and found only DENV counts at the level of Myanmar’s states for the year 2015 as reported by the Ministry of Health and Sports [30]. Similarly to the approach applied for districts, we calculated the yearly average index P per state using climate data for both 2015 and 2016 (Figure 2B) and compared its spatial distribution with DENV counts per state for 2015 (Figure 2C). States presenting higher average suitability were located across the centre of the country, but particularly in the south, sharing a border with Thailand. The distribution of DENV counts presented a generally similar spatial signature co-localised in the centre and south of the country (Figure 2C). The Pearson’s correlation between mean yearly suitability and yearly DENV counts of the state-based two maps was 0.56 (p-value=0.033).

Again, since we propose the index P as a measure of suitability independent of the number of (female) mosquitoes per human (M), a typical threshold of P=1 may not be adequate to speculate on local epidemic potential, given that an R0>1 can be achieved with P<1 and M>1 (although the results from Figure 1 suggest that both the timing and amplitude of P are highly informative). We hypothesized that the time spent above a certain value (threshold) of suitability could be a better proxy for local, yearly epidemic potential. A sensitivity analysis was performed for 2015, the only year for which we had both climate input (index P) and DENV counts, with the intent of searching for the threshold of P that best explained the observed spatial distribution of DENV counts per state. We set 100 thresholds from 0.5 to 2.5 and measured the amount of time (T) the index P remained above each threshold in each state in 2015. Measures of T per state were used to calculate Pearson’s correlation with DENV counts per state (Figure 2D). We found that small thresholds (P<1) had non-significant and low correlations between yearly cases and mean yearly suitability, while intermediate-to-high thresholds (P>1.35) had significant and high correlations. The minimum correlation was found Just above P=1 (dashed vertical lines in Figure 2D), when states are seen to spend equal amount of time T above that threshold in a year (seen by measures of T per state coloured according to DENV counts as in map’s legend, Figure 2D). These results show the amount of time a region spends with suitability above a typical threshold of 1 is the least optimal to predict yearly epidemic potential and that in Myanmar’s epidemiological context higher thresholds are more informative.

## Discussion

In this study, we were able to demonstrate that our measure for mosquito-borne viral suitability is informative in the context of Myanmar, despite the lack of ento-epidemiological datasets with high spatio-temporal resolution. By estimating suitability through climate variables and known ento-epidemiological parameters, we were able to project mosquito-born virus (MBV) suitability at the district level, a resolution for which epidemiological data and mosquito abundance measures are not generally available. Here, we discuss the national and subnational public health and control implications for MBVs in the context of our projections.

At both the national and subnational levels, the wet season (Jun - Oct) was estimated to have the highest potential for MBVs transmission, in accordance with reported epidemiological time series. Since this was observed across Myanmar, it suggests that the epidemic potential of each district peaks during this period independently of their spatial location. In contrast, the hot and dry season (Mar - May) presented the lowest potential, also consistently across Myanmar, in accordance with what are known to be less favourable climate conditions for the vector. We therefore argue that in Myanmar, adequate surveillance and health care delivery resources should be fully operational by the end of the hot and dry season (May), in anticipation for the increase in MBVs case counts that is likely to occur in the following months. A similar argument can be used for vector control strategies in Myanmar, which should have maximal impact before the onset of the wet season, when mosquitoes encounter less favourable climate conditions and have smaller population sizes.

We also identified important spatial variations in MBV suitability across Myanmar. For instance, the highest potential was found primarily in the southern rural districts bordering with Thailand, and to a lesser degree across the middle of the country where the 3 major urban centres are located. Importantly, this estimated potential was highly correlated with DENV cases counts at the state level, although the lack of epidemiological data at such spatial resolution precluded us from verifying if our estimations at the district level fitted local counts. It may be tempting to speculate that the higher number of DENV counts in the south is a consequence of inflow of cases from Thailand (a highly endemic country). Critically, however, the estimated suitability in the south of Myanmar, in particular during the wet season, suggests for the first time that the observed higher incidence is driven by a local, higher intrinsic potential for MBVs transmission. Although management of case importation should be part of any national plan, it is clear that reducing suitability in the southern region of the country will be critical to control MBVs, for which vector control strategies should be effectve.

Epidemiological models are useful tools to gauge the burden of a disease of interest, assess transmission potential, mosquito suitability, prompt surveillance efforts, inform better public health policies and highlight areas for pressing research. Such approaches are even more critical in epidemiological settings characterized by the absence of sustained surveillance or for pathogens which tend to have mild or asymptomatic pathology. The method here introduced requires solely climatic data and basic ento-epidemiological assumptions for which literature support is available. We foresee the usefulness and applicability of the index P for other regions of the world for which surveillance data is still missing, either due to lack of resources or absence of a pathogen of interest. In contrast, for regions rich in historical spatio-temporal and ento-epidemiological data, the index P may be a starting point for the development of an early warning system which would be based on real-time input of climatic variables.

## Implications for Zika virus in Myanmar

Our understanding of Zika virus (ZIKV) epidemiology in Myanmar and other countries of South East Asia (SEA) is incomplete, although there is evidence of continued transmission in the region from serosurveys and occasional viral isolation in residents and travelers to the region [31]–[34]. Such evidence also supports the notion that ZIKV transmission in SEA preceded that of the Americas (2014-2015, [35]) even with an apparently low number of cases and no major epidemics reported. To date, only one imported ZIKV case has been notified by the Myanmar Ministry of Health and Sports (MOHS) [36] and the virus’ spatio-temporal potential for transmission in the country is largely unknown. Given that mainland countries of SEA share many of the climatic and eco-demographic factors that dictate positive suitability for *Aedes* mosquitoes, it is reasonable to assume that regions within Myanmar have the potential for epidemic or endemic transmission of ZIKV. Exploiting the fact that transmission seasons of various Aedes-born viruses (e.g. DENV, CHIKV, ZIKV) tend to be synced in time in other regions of the world [37], we here discuss and speculate on the ZIKV public health implications of the index P’s spatio-temporal patterns found both at the national and subnational levels in Myanmar.

The higher suitability for MBVs in southern Myanmar suggests the south to be a viable route for importation of ZIKV from Thailand. It is known that southern international borders are home to sizeable mobile populations with limited access to healthcare [38]. Introduction of ZIKV through such borders would therefore carry a significant public health burden but would also likely be difficult to detect with a passive surveillance system. Additionally, suitability and DENV counts suggest a path of high transmission potential in the middle of the country, in districts including the 3 major urban centres of Myanmar (Yangon, Naypyitaw and Mandalay). Due to the domestic nature of the mosquito species involved, urban centres are a hallmark for ZIKV transmission and establishment, with attack rates above 60% reported elsewhere [15], [39], [40]. For the city of Yangon, for example, a similar attack rate would result in +3 million cases, and would incur significant health and economic consequences. Public health prevention or mitigation of a starting epidemic therefore calls for active surveillance initiatives that move beyond formal international points of entry (i.e. airports and maritime ports) and urban centres. Detecting early epidemic transmission chains in time for mosquito-control interventions before the wet season may effectively hamper the full potential of ZIKV and prevent high attack rates.

Exposure to ZIKV infection during gestation is a major risk factor for development of a variety of neonate neurological complications including microcephaly (MC) [35], [41], [42]. Recent studies have further suggested that the risk of MC is highest for exposure around week 17 of gestation, resulting in a lag of approximately 5 months between ZIKV and MC epidemic peaks [15], [41]–[43]. Based on the estimated time window of peak MBV suitability in Myanmar between June and October, we therefore predict that, in the event of a ZIKV outbreak, an epidemic of MC in the country would occur between November and March. This time window is therefore critical for active MC surveillance to be established in Myanmar. To date, there has been no report of significant increases in MC cases in Myanmar. Caution should be taken in assuming this as evidence for no ZIKV circulation, since in previous epidemics there has also been a lack of reported ZIKV-associated MC cases [44] and it is possible that only one of two existing lineages of the virus is responsible or such clinical manifestations [35], [41], [42], [45].

Put together, the estimated spatio-temporal variations in MBV suitability found in this study suggest that in order to decrease mosquito populations before the onset of ZIKV epidemics or prevent potential ZIKV introduction events from the southern region, control initiatives should take place just before, and at the beginning of, the wet season. Special attention could also potentially be stratified across districts or states in the middle of the country, including the major urban centres, and in particular in the south, as these regions are likely to have higher transmission potential.

## Limitations and future work

There are certain limitations to our approach. We note that the unavailability of high resolution climatic data meant that (1) it was impossible to estimate suitability along the border with China and Laos, two countries in which DENV transmission is reported to be endemic; and (2) that weather stations had to be used for vast geographical ranges, limiting our capacity to explore potentially relevant spatial heterogeneities within the larger districts. The climatic data used was also limited to 2 years, and although we show that the index P in that period explains much of DENV’s epidemiology in 1996-2001, it is uncertain to what degree our estimations could have been better with matching time periods. It should also be noted that we take care in not interpreting P>1 as a critical threshold for transmission potential. The real epidemic thresholds (R0>1, Re>1) are dependent on the total number of female mosquitoes per human (M=NV/NH) which is largely unknown in time and space. In this context, our sensitivity analysis in Figure 2D helps to elucidate this and can be of use for other regions for which climatic variables are available. Another climatic data source that could be investigated in future is satellite remote sensing, although we did not use this for the present study. We also discuss the implications for ZIKV transmission in Myanmar, although our results are based on DENV epidemiological data. Given that no seroprevalence or epidemiological data exists for ZIKV in Myanmar, our discussion points are intended to inform the community to the best of our knowledge, but should be taken as speculative until new data is made available and compared to our current projections. Our approach also does not include demographic factors, why may affect both the human susceptibility and the vector carrying capacity. Although our index P can explain much of the spatio-temporal patterns of Myanmar, it is possible that these factors explain some of the missed spatial patterns (e.g. local vector capacity could be higher in regions with more cases than predicted by P).

## Ento-epidemiological count data

DENV case counts for Myanmar between 1996 and 2001 (as published by [12]) were published already aggregated at the level of the country and by month. Naing et al. reported that the original source of the case counts was the official annual reports of the Myanmar National Vector-Borne Disease Control programme (VBDC). Cases included total suspected reports of dengue fever (DF) and dengue haemorrhagic fever (DHF). The absolute counts, per month (Jan-Dec) were: 20, 35, 40, 43, 139, 333, 487, 255, 258, 261, 80, 82 (year 1996); 67, 68, 37, 55, 140, 724, 876, 877, 545, 430, 248, 150 (year 1997); 131, 144, 208, 271, 714, 2511, 2904, 2455, 1528, 1163, 747, 249 (year 1998); 45, 105, 164, 91, 445, 1309, 1533, 968, 533, 247, 92, 102 (year 1999); 32, 29, 26, 90, 120, 333, 164, 95, 59, 73, 71, 84 (year 2000); 16, 29, 576, 1190, 2137, 2868, 3082, 2346, 1476, 990, 354, 51 (year 2001).

DENV counts at the level of Myanmar’s states for the year 2015 were used, as reported by the Ministry of Health and Sports [30]. Incidence per 100k individuals per state were: Sagaing 125, Ayeyarwady 105, Mandalay 106, Mon 259, Yangon 63, Bago 68, Tanintharyi 145, Naypyitaw 163, Shan 50, Kayin 108, Magway 35, Kachin 61, Rakhine 28, Kayah 102, Chin 18.

Data collected in a study of *Aedes aegypt* abundance in Yangon over the year of 2011 was also used [29] From this publication we used the number of major breeding containers found in Yangon per month (Jan-Dec): 110, 102, 58, 51, 109, 249, 257, 276, 258, 248, 252, 170.

## Spatial and climatic data

The administrative distribution of Myanmar into districts was suitable for our analysis, since it was possible to classify them by predominant weather conditions, using the Köppen-Geiger classification [46]: equatorial monsoonal (Am), equatorial winter dry (Aw), warm temperate-winter dry-hot summer (Cwa), warm temperate-winter dry-warm summer (Cwb) and arid steppe-hot arid (BSh). We obtained climate data from the United States National Oceanic and Atmospheric Administration webpage [47], which had incomplete observations that we then complemented with information from the Department of Meteorology and Hydrology, Yangon, for the period 2015-2016. Time and resource constrains for this process of data collection allowed for retrieving data from 14 weather stations, which were representative of the following districts: Pathein station, for the districts of Pathein, Pyapon, Maubin, Myaungmya and Labutta; Hpa An station, for Hpa An, Myawaddy, Kawkareik, Mawlamyine and Thaton; Sittwe station, for Sittwe, Marauk-U and Maungdaw; Dawei station, for Dawei, Myeik, and Kawthoung; Yangon Airport station, for North, South, East and West Yangon; Bago station for Bago, Hpapun and Hinthada; Nay Pyi Taw Airport station for North and South Nay Pyi Taw, Yamethin and Magway; Loikaw station for Loikaw, Bawlake and Langkho; Katha station for Katha, Bhamo and Mohnyin; Hkamti station for Hkamti district only; Taunggyi station for Taunggyi and Loilen; Mandalay Airport station for Mandalay, Kyaukse, Miyngyan, Nyaung-U and Meiktila; an average of the weather conditions in the Am region, for the districts of Kyaukpyu and Thandwe); and an average of the weather conditions in the Aw region, for the districts of Minbu, Pakokku, Gangaw, Pyinoolwin, Sagaing, Shwebo, Monywa, Kale, Yinmabin and Kyaukme.

To include a district in the present analysis, we used the criteria that its main population settlements were below 1500 meters above sea level, since the entomological modelling system we employed does not account for the effect of elevation on vector ecology and higher altitudes are less favourable to mosquito survival, plus either of the following: having access to climate variables from its weather station; or that its central point was within 100 Km of a station from which climate information was available; or being situated within a weather region were climate could be extrapolated from other districts’ stations. The latter was done since an analysis of variance showed no difference in mean temperature across weather stations in the Am (F=0.391, p-value=0.53; Dawei, Hpa An, Yangon, Pathein and Sittwe, and Bago stations) and Aw regions (F=2.793, p-value=0.09; Taungoo, Loikaw and Nay Pyi Taw stations). Extrapolation was not done for districts within the Cwa region, as there was a statistically significant difference in weather observations from individual stations (F=12.03, p-value<0.05; Hkamti, Katha nad Taunggyi stations). Lastly, Mandalay was a single station within the BSh region and Hakha station from the Cwb region was removed from analysis, due to elevation criteria.

The three weather seasons defined in this study for the context of Myanmar were: cool dry season, from November to February, hot dry season, from March to May, and wet (monsoon) season, from June to October. National means (standard deviations) of yearly and cool, hot and wet season were, correspondingly: temperature in degrees Celsius 27.1 (2.8), 26.7 (1.5), 28.8 (1.3), and 27.7 (1.4); percent humidity 77.9 (11.1), 79.9 (5.3), 64.5 (5.8), 85.3 (6.7); and inches of rainfall 0.17 (0.43), 0.03 (0.13), 0.01 (0.05) and 0.47 (0.62).

## Parameters specific to Myanmar

As detailed in the main text, unknown parameters α, ρ and η were obtained for each weather station using a parameter sweep with heuristics of adult mosquito life-span of ~9 days and extrinsic incubation period of ~ 5 days (over the period 2015-2016). See Methods for details. The obtained values for α, ρ and η per weather station were (in order): Pathein 1.414, 0.78, 2.241; Hpa An 1.414, 0.78, 2.241; Sittwe 1.552, 0.45, 2.517; Dawei 1.414, 0.56, 2.517; Yangon 1.552, 0.78, 2.241; Am 1.414, 0.78, 2.379; Bago 1.414, 0.67, 2.379; Nay Pyi Taw 1.552, 0.45, 2.517; Loikaw 1.966, 0.45, 2.379; Aw 1.552, 0.78, 2.241; Katha 1.689, 0.56, 2.379; Hkamti 1.828, 1.00, 1.828; Taunggyi 1.828, 0.78, 1.828; Mandalay 1.552, 0.23, 2.655.

The districts found to have yearly mean index P>1 were: Magway, Nay Pyi Taw north, Nay Pyi Taw south, Yamethin, Bhamo, Katha, Mohnyin, Dawei, Kawthoung, Myeik, Maungdaw, Mrauk-U, Sittwe, Bago, Hinthada, Hpapun, Hkamti, Hpa-An, Kawkareik, Mawlamyine, Myawaddy, Thaton (with no particular order).

